# AlphaFold3, a secret sauce for predicting mutational effects on protein-protein interactions

**DOI:** 10.1101/2024.05.25.595871

**Authors:** Wei Lu, Jixian Zhang, Jiahua Rao, Zhongyue Zhang, Shuangjia Zheng

## Abstract

AlphaFold3 has set the new state-of-the-art in predicting protein-protein complex structures. However, the complete picture of biomolecular interactions cannot be fully captured by static structures alone. In the field of protein engineering and antibody discovery, the connection from structure to function is often mediated by binding energy. This work benchmarks AlphaFold3 against SKEMPI, a commonly used binding energy dataset. We demonstrate that AlphaFold3 learns unique information and synergizes with force field, profile-based, and other deep learning methods in predicting the mutational effects on protein-protein interactions. We hypothesize that AlphaFold3 captures a more global effect of mutations by learning a smoother energy landscape, but it lacks the modeling of full atomic details that are better addressed by force field methods, which possess a more rugged energy landscape. Integrating both approaches could be an interesting future direction. All of our benchmark results are openly available at https://github.com/luwei0917/AlphaFold3_PPI.

## 1 Introduction

The development of AlphaFold and other deep learning methods has revolutionized the study of the protein complex structures [1–7], advancing beyond traditional physics-based docking methods [8–10]. With the advent of AlphaFold3, the success rate of predicting general protein complex structures has reached almost 80%, and that for protein-antibody pairs has exceeded 60%, significantly outperforming its predecessor, AF2-Multimer, previously considered the state-of-the-art [1, 2]. However, as Derek Lowe points out [11], “Structure is not everything.” The complete picture of biomolecular interactions cannot be fully captured by static structures alone; it also involves the dynamic association and dissociation of one protein with another protein partner. In an equilibrium state, this interaction property is commonly measured as the binding affinity, *K*_*d*_, or described as the change in binding free energy, Δ*G*. In addition, nature is constantly evolving, with mutations regularly introduced that modulate the magnitude of binding affinity, among many other properties, such as the stability of the proteins themselves. A key therapeutic goal is to design molecules that bind strongly enough to either prevent or promote a specific state of its binding partner, such as the inactive or active state of GPCRs. In the field of antibody discovery, the process of mutating a candidate antibody to improve its binding affinity with the target antigen protein is termed antibody maturation. Most antibody drugs in clinical use have picomolar affinity for the target protein [12], while antibodies found through immunization commonly exhibit affinities in the nanomolar or even micromolar range [13]. This means that, during the maturation step, we need to improve the affinity by more than a thousand times, or, if measured in ΔΔ*G*, by about 4 kcal/mol.

A variety of methods have been developed to estimate the effects of mutations, ranging from force field-based methods [14, 15], which derive forces from physical interactions such as van der Waals and electrostatic, or from statistical energy, to profile-based methods [16], which query sequence and structure databases. Additionally, there are hybrid methods [17] that combine force field and profile-based information. More recently, deep learning methods [18–21] have emerged, where the underlying energy landscapes are learned through unsupervised pre-training involving perturbations of crystallized protein structures.

In this work, we demonstrate that AlphaFold, although trained as a structure prediction model, learns critical information that is complementary to all types of current methods studying mutational effects on protein-protein interactions. This observation aligns with findings from previous studies of protein-small molecule and protein-peptide interactions [22–25], suggesting that models also trained to predict complex structures demonstrate enhanced capabilities in predicting binding affinity.

## 2 Related Work

Many works have demonstrated that AlphaFold2 already learns important features useful for other tasks. For example, in the protein design field, RFdiffusion [26] showed that fine-tuning from the pre-trained structure prediction model, RoseTTAFold [4], significantly enhances performance in protein design compared to starting without pre-training. Additionally, Roney [27] demonstrated that AlphaFold2 discerns the underlying physics capable of differentiating decoy structures from native structures, thereby effectively ranking candidate complex structures.

Other works utilize AF2 outputs directly as inputs for their models. Akdel [28] demonstrated that the AF2-predicted structure could serve as the input for many popular structure-based predictors of protein thermostability, such as FoldX and DynaMut2 [14, 29], achieving results comparable to those obtained with crystal structures. Additionally, Lyu [30] showed that AlphaFold2 structures could be used as inputs for docking programs in small molecule drug discovery. Although Buel [31] showed that AlphaFold2 cannot predict key mutational effects in many cases, McBride [32] demonstrated that with appropriately chosen metrics, such as effective strain in their study, AlphaFold2 can accurately predict the effects of mutations on the intrinsic properties of single proteins, using three experimental datasets: fluorescence, folding, and catalysis.

These works have explored the utility of AlphaFold in many research areas, but none of them have studied the usefulness of AlphaFold in predicting the binding energy and the mutational effects of mutations on protein-protein interactions.

## 3 Benchmark setups

A commonly used dataset for evaluating methods that predict mutation effects is SKEMPI [33], which manually curates a list of crystallized protein complexes and their mutants with experimentally measured changes in binding affinity, denoted as ΔΔ*G* values, gathered through literature searches. These values are measured using biochemical methods that, while relatively accurate, require considerable effort for each mutant’s data. As a result, despite the intensive labor involved, this dataset contains binding data for only 7085 mutations, which is significantly fewer than the total number of crystallized structures, and even less than the number of sequences in the database that enabled the development of AlphaFold. This scarcity of data underscores the value of leveraging information learned from tasks beyond direct protein-protein binding data.

### 3.1 Dataset definition

In order to benchmark against a wide range of methods, we utilized the common subset of Test Set 1 from SKEMPI as defined in SSIPe [17] and the SKEMPI dataset as employed in DSMBind [19]. The cases where the ranking score predicted by AlphaFold3 is below 0.8 are removed. As a result, our benchmark comprises 475 mutants across 42 unique protein complexes.

### 3.2 Baselines

We include a comprehensive set of 17 baseline methods for benchmarking. These can be categorized by their types of approaches.

#### Protein Language-based Models

ESM2, ESM1v, and ProGen2 [34–36] are prominent protein language models known for their robust zero-shot performance across multiple tasks, including secondary structure prediction, and the classification of benign and pathogenic mutations. Given that these models typically accept only a single sequence as input, we concatenated the sequences of interacting proteins for our analysis.

#### Force Field and Profile-based Models

Included in our baselines are three popular pure force field-based models: FoldX, FlexddG, and EvoEF [14, 15, 37]; a structure-based profiling baseline, BindProfX [16]; and one hybrid model that combines sequence and structure profiling with force field methods, SSIPe [17]. These models are specifically designed to predict the mutational effects on protein-protein interactions, with SSIPe considered the state-of-the-art model that utilizes the most information available.

#### Structure-based Deep Learning Models

ProteinMPNN [38] is a model that learns to design sequences corresponding to a given input backbone structure, while DSMBind [19] predicts mutational effects by learning to restore a perturbed crystal structure. Both models develop a scoring function that can be used to estimate mutational effects for a given input structure and corresponding sequences.

#### AlphaFold3, AlphaFold2, and Strain

Protein sequences are submitted directly to the AlphaFold3 server in JSON format with the seed set to a fixed arbitrary number, 42, to ensure reproducibility. Five results are downloaded, and the top one is used. For each mutant, the predicted score is calculated by subtracting the ranking score of the wild type from its ranking score. AF2-Multimer v2.3.0 [2] is run locally with default settings, and the ranking score for the top predicted model for each entry is used. As a simple statistical baseline, effective strain, as defined in [27], is computed for each mutant.

## 4 Results

### 4.1 Comparison of binding energy estimation across all baselines

The results for each baseline are summarized in Table 1. Pearson and Spearman correlation coefficients are two commonly used metrics for assessing continuous variables. The Area under the ROC Curve (AUC) is a statistic used in binary classification, computed by treating all ΔΔ*G* values below zero as positive and those above zero as negative. The results are sorted according to their Pearson correlation coefficients within each category.

**Table 1:** Comparison of ΔΔ*G* estimation results on our SKEMPI test set using three different metrics.

| Category | Method | Pearson | Spearman | AUC |
| --- | --- | --- | --- | --- |
| Force Field and Profile-based | SSIPe | 0.68 | 0.62 | 0.78 |
|  | FlexddG | 0.62 | 0.58 | 0.77 |
|  | BindProfX | 0.56 | 0.58 | 0.74 |
|  | EvoEF | 0.55 | 0.51 | 0.72 |
|  | FoldX | 0.49 | 0.54 | 0.74 |
| Structure-based Deep Learning | DSMBind | 0.62 | 0.53 | 0.73 |
|  | ProteinMPNN | 0.51 | 0.45 | 0.65 |
| AlphaFold | AF3 ranking_score | 0.49 | 0.51 | 0.71 |
|  | AF3 iptm | 0.49 | 0.50 | 0.72 |
|  | AF3 ptm | 0.36 | 0.33 | 0.63 |
|  | AF3 mean_pae | 0.32 | 0.37 | 0.64 |
|  | AF2 ranking_score | 0.21 | 0.23 | 0.57 |
|  | Effective Strain | 0.18 | 0.31 | 0.61 |
|  | AF2 mean_pae | 0.05 | 0.22 | 0.54 |
| Protein Language-based | ESM2 | 0.27 | 0.35 | 0.68 |
|  | ESM1v | -0.02 | 0.06 | 0.52 |
|  | ProGen2 | -0.09 | 0.01 | 0.47 |

Table 1 indicates that protein language models are less effective at predicting the mutational effects on protein-protein interactions. Similarly, AlphaFold2 and Strain show weak correlations with experimentally measured binding affinities. In contrast, the ranking score produced by AlphaFold3 exhibits a significant correlation and is comparable to that of the widely-used FoldX method.

### 4.2 AlphaFold3 complements other baselines

As demonstrated by SSIPe [17], profile-based and force field-based methods complement each other, producing the most accurate estimators. Similarly, if AlphaFold3 learns a different type of information, it could also complement other models. As shown in Fig 1, a simple ensemble of AlphaFold3’s ranking scores boosts performance across all baselines. The ensemble score is computed by adding the equally weighted ranked scores of two models. Notably, the previous state-of-the-art, SSIPe, which is already a combination of models, also experiences a performance boost. AlphaFold3 learns complementary information that is orthogonal to current methods, thereby enhancing the estimation of mutation effects on protein-protein interactions.

**Figure 1:**
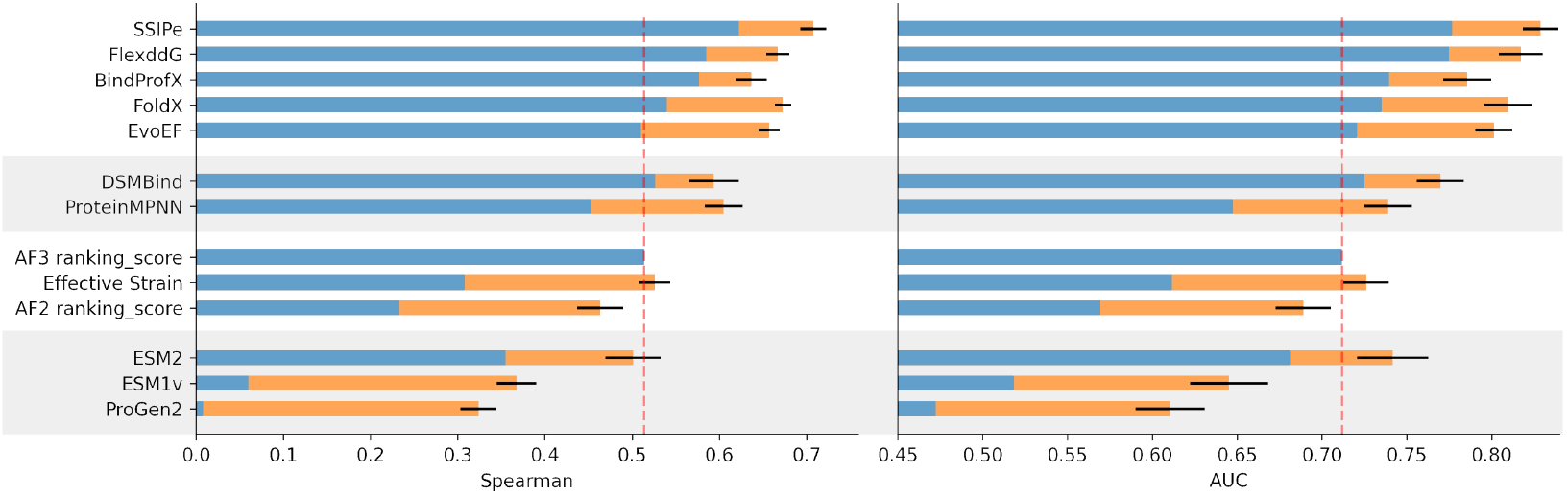
Ensemble with AlphaFold3 boosts performance across all baselines, as evaluated by Spearman correlation, **Left**, and AUC, **Right**, blue is the baseline score, orange is the boost in performance after ensemble with AlphaFold3 score. The dashed line indicates the AlphaFold3 performance.

### 4.3 AlphaFold3 offers unique information

To investigate whether the predictions made by AlphaFold3 are correlated with those of other methods, we computed the pairwise correlation among all methods, as shown on the left of Fig. 2. AlphaFold3 exhibits very weak correlations with other models, only showing slight correlation with DSMBind. In contrast, other models, such as FlexddG and SSIPe, correlate with many other methods, indicating that AlphaFold3 learns unique features that are orthogonal to those of other methods. As shown on the right side of Fig. 2, protein language models, AlphaFold2, and strain do not provide additional information beyond what AlphaFold3 provides. Conversely, structure-based deep learning, as well as force field and profile-based methods, enhance the predictions made by AlphaFold3.

**Figure 2:**
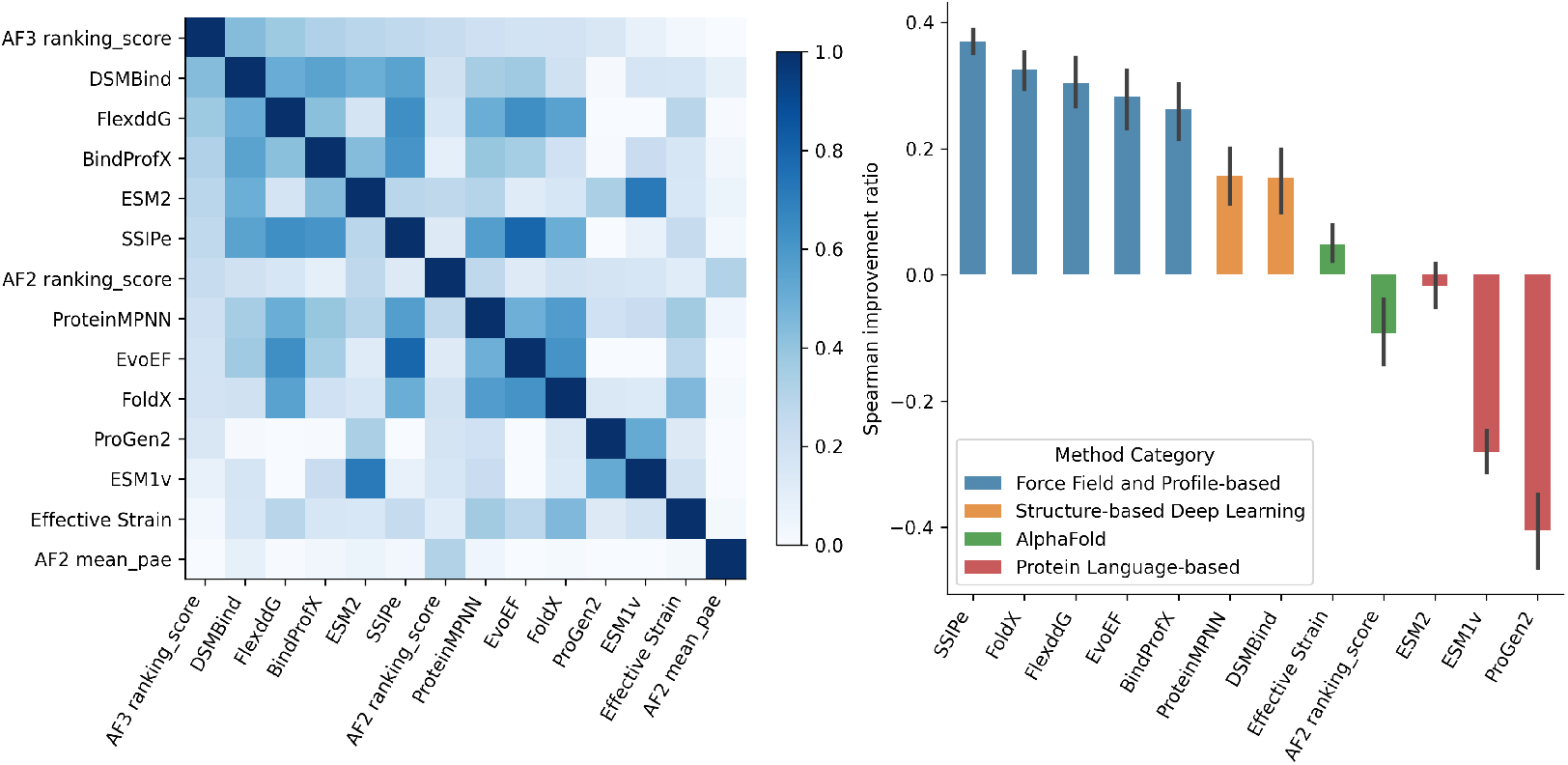
**Left**, correlation between model predictions, sorted by their correlation against AlphaFold3. **Right**, when AlphaFold3 is combined with other models, all methods except protein language models and AlphaFold2 get a significant boost. y axis measures the ratio of improvement relative to the AlphaFold3 baseline, (*r*_*m*+*AF*3_ − *r*_*AF*3_)*/r*_*AF*3_

## 5 Discussion

Our results indicate that the performance of protein language models in predicting the mutational effects on binding affinity is relatively weak. This finding aligns with a recent study [39], which demonstrates that language models do not scale effectively with model size in prediction tasks that are less dependent on coevolutionary patterns.

An interesting distinction between AlphaFold3 and traditional all-atom force field methods lies in their sensitivity to protein complex conformations. Traditional force fields are highly sensitive to the exact conformation of the protein complex, whereas AlphaFold3 tends to predict similar scores for identical input sequences. This sensitivity to conformation and the difficulty in thoroughly sampling conformations make force field methods prone to inaccurate estimations of the entropy component of the total Gibbs free energy. In contrast, AlphaFold3, as a generative structure prediction model, is capable of learning a smoother energy landscape that more effectively captures the subtle influences of entropy. Fig 3 illustrates how the integration of AlphaFold3 scores can more accurately determine the global relative free energy, Δ*G*, compared to relying solely on force field methods, which exhibit a more rugged energy landscape.

**Figure 3:**
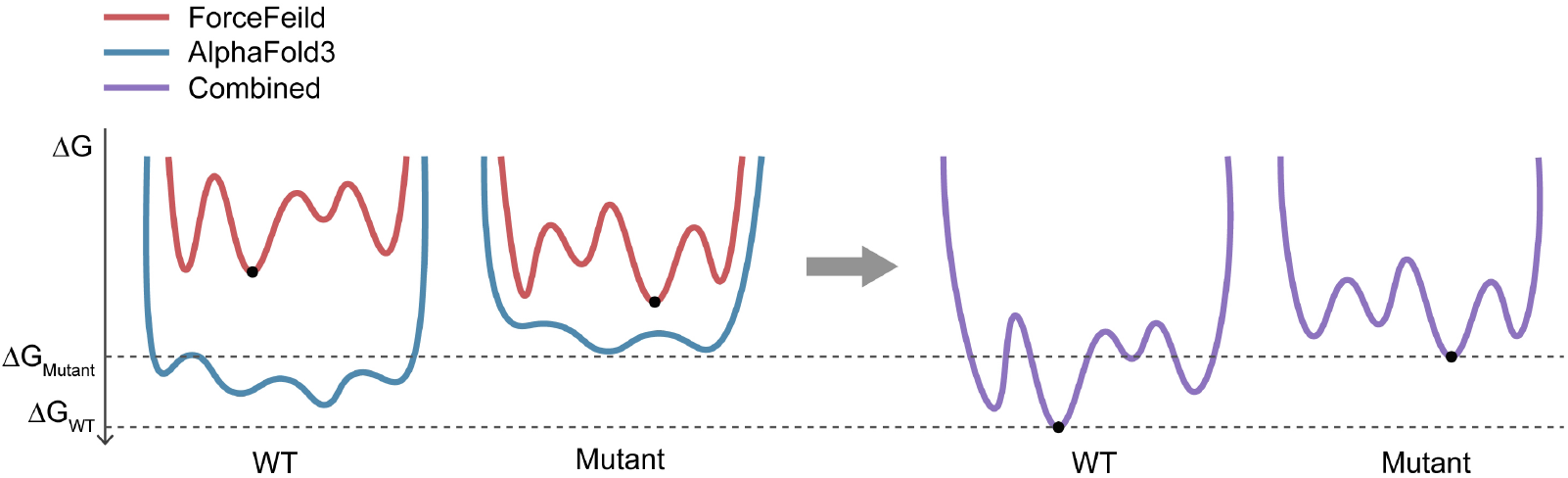
A schematic plot illustrating how the rugged energy landscapes of all-atom force fields (in red) when combined with the globally more accurate energy landscape of AlphaFold3 (in blue), result in a composite energy landscape (in purple) that more closely approximates the ground truth (dashed lines) for estimating Δ*G*.

## 6 Conclusion

In this study, we have demonstrated that AlphaFold3 learns unique features beneficial for estimating binding free energy and complements existing models. Looking ahead, a more integrated approach combining folding methods that predict complex structures with inverse-folding models that identify masked residues, along with traditional force field and profile-based models, could significantly revolutionize the field of predicting mutational effects on protein-protein interactions.

## Acknowledgments and Disclosure of Funding

We thank our colleagues for submitting the test queries to the AlphaFold3 server. The authors declare that there are no competing interests associated with this work. This work was partly supported by Young Elite Scientists Sponsorship Program by CAST and RayWu Angel Fund.

